# Imaging the extent and location of spatiotemporally distributed epileptiform sources from MEG measurements

**DOI:** 10.1101/2021.11.09.467915

**Authors:** Xiyuan Jiang, Shuai Ye, Abbas Sohrabpour, Anto Bagić, Bin He

## Abstract

Non-invasive MEG/EEG source imaging provides valuable information about the epileptogenic brain areas which can be used to aid presurgical planning in focal epilepsy patients suffering from drug-resistant seizures. However, the source extent estimation for electrophysiological source imaging remains to be a challenge and is usually largely dependent on subjective choice. Our recently developed algorithm, fast spatiotemporal iteratively reweighted edge sparsity minimization (FAST-IRES) strategy, has been shown to objectively estimate extended sources from EEG recording, while it has not been applied to MEG recordings. In this work, through extensive numerical experiments and real data analysis in a group of focal drug-resistant epilepsy patients’ interictal spikes, we demonstrated the ability of FAST-IRES algorithm to image the location and extent of underlying epilepsy sources from MEG measurements. Our results indicate the merits of FAST-IRES in imaging the location and extent of epilepsy sources for pre-surgical evaluation from MEG measurements.

## Introduction

Epilepsy is one of the major neurological diseases affecting more than 65 millions of people globally (Moshé et al., 2015; Thijs et al., 2019). In about 30% of these patients, seizures cannot be controlled with medications alone, and brain surgery can be a viable treatment option for a selection of these patients (Choi et al., 2008; Lamberink et al., 2017), if the epileptogenic zone can be accurately localized and safely removed. Thus, a successful pre-surgical evaluation is needed to determine the epileptogenic zone (EZ), which is the minimum amount of cortex that needs to be resected to achieve seizure freedom (Duncan et al., 2016; Dwivedi et al., 2017; Rosenow and Lüders, 2001). Currently, the gold standard for pre-surgical evaluation is the intracranial electroencephalography (iEEG) (Wilke et al., 2011; Yuan et al., 2012; Dichter, 2014; Jayakar et al., 2016; Jiang et al., 2019), where electrodes are placed over the presumed epileptogenic regions to record epileptogenic brain activity such as the interictal discharges and ictal activity. Nevertheless, limited by its invasive nature, iEEG recording could not cover a large cortical area and adds risks to the patients (Arya et al., 2013; Malmgren and Edelvik, 2017).

On the other hand, non-invasive electromagnetic recording techniques such as MEG and EEG record the whole-brain activity, and are also suitable for the pre-surgical evaluation of epilepsy patients (Luders, 2008; He et al., 2018, 2020; Bagic and Burgess, 2020). EEG and MEG are able to resolve neural activity with high temporal resolution, in the order of milliseconds (Hämäläinen et al., 1993; da Silva, 2013; He et al., 2018), but the spatial resolution is limited and needs to be further boosted (Baillet, 2017; Michel and He, 2017; He et al., 2018). Electrophysiological source imaging (ESI) is one such effort to improve the spatial resolution of EEG and MEG. In short, ESI estimates the underlying brain electrical activity from EEG/MEG measurements, by modeling these electrical activities as equivalent current dipoles or distributed current sources (Michel et al., 2004; He et al., 2018).

Various biomarkers have been analyzed by means of ESI techniques to delineate epilepsy networks subtending seizures (Yang et al., 2011; Lu et al., 2012; Sohrabpour et al., 2020; Ye et al., 2021), high-frequency oscillations (HFOs) (Lu et al., 2014; Tamilia et al., 2017; Thomschewski et al., 2019; Velmurugan et al., 2019; Cai et al., 2021) and interictal spikes (Wang et al., 2011; Sohrabpour et al., 2016b, 2020; Ye et al., 2021). Interictal spikes, which are brief electrographic discharges that occur in between seizures (de Curtis and Avanzini, 2001), are of special importance when studying epilepsy and are widely used as a biomarker to approximate the EZ from EEG/MEG data (Ebersole, 1997; Mégevand et al., 2014; Englot et al., 2015; Ye et al., 2021).

ESI algorithms are constantly evolving (Van Veen et al., 1997; Pascual-Marqui, 2002; Grova et al., 2006; Ding and He, 2006; Wipf and Nagarajan, 2009; Zhu et al., 2014; Chowdhury et al., 2016; Krishnaswamy et al., 2017; Cai et al., 2018), but a common shortcoming for most of the previous ESI algorithms is that they do not give an explicit estimation of the extent of the underlying sources (Sohrabpour et al., 2016a; Sohrabpour and He, 2021). Determining underlying sources’ extent is of significance and highly desirable, as different brain regions are specialized to perform different functions and it is desirable to image the location and extent of these functional areas, noninvasively. Additionally, when imaging pathological activities such as epileptic networks, not being able to estimate the underlying sources’ extent would deprive us from knowing and precisely delineating the epileptogenic brain tissues, as important information about the extent of the pathological region is missed (Vakharia et al., 2018).

One of the recent developments in filling this gap is a new framework called, “fast spatiotemporal iteratively reweighted edge sparsity minimization (FAST-IRES)” algorithm (Sohrabpour et al., 2020), which objectively estimates extended sources and their time-course of activity by solving a series of convex optimization problems. Specifically, it has been shown that FAST-IRES is capable of estimating the location and extent of underlying brains sources from noninvasive scalp EEG recordings, in synthetic (simulated) and real data.

While EEG and MEG are electromagnetic field recordings of brain activities and bear a lot of similarities (Iwasaki et al., 2005; Knake et al., 2006), they measure, in principle, different physical quantities, and have different sensitivity with respect to source properties (Baumgartner, 2004; Baillet, 2017). While EEG is impacted by the low conductivity of the skull (Tuch et al., 2001; Vorwerk et al., 2014; Dabek et al., 2016; Fiederer et al., 2016), MEG is blind to radial sources due to the sensitivity of MEG sensors (Ahlfors et al., 2010; Piastra et al., 2021). MEG offers various types of sensors, such as the magnetometer which measures the strength of the magnetic field (MAG), and the gradiometer which measures the spatial gradient of the magnetic field (GRAD) (Wynn et al., 1975; Malmivuo et al., 1995). The MEG community is still divided in terms of which configuration should be used when doing source reconstruction, including prior work that uses MAG only (Assaf et al., 2004; García-Pacios et al., 2015), GRAD only (Popescu et al., 2010; Wens et al., 2014), and combining MAG and GRAD (Henson et al., 2009; Hillebrand et al., 2016). In principle, the combination of MAG and GRAD emphasizes brain signals with respect to environmental noise (Kominis et al., 2003; Baillet, 2017), but there is prior work suggesting that no significant improvement is achieved by combining MAG and GRAD sensors for the purposes of MEG source imaging (Tarkiainen et al., 2003; Garcés et al., 2017). Given the current debate, it is of interest to see how FAST-IRES performs using signals recorded from these different sensor group configurations.

The source extent estimation in distributed MEG source imaging remains to be a challenge and is on its early stage. The most common way is to set a threshold for the solution distribution, for instance based on empirical data (Attal and Schwartz, 2013; Pellegrino et al., 2018); the choice of the threshold, however, is highly dependent on the researcher’s experience. There is also prior work using post-hoc analysis such as receiver operative curves (Bouet et al., 2012), or maximum beamformer output (Hillebrand and Barnes, 2011), but these methods have limitations where they require prior knowledge of the ground truth (Hillebrand and Barnes, 2011; Bouet et al., 2012), which is not available in realistic applications. Other studies have proposed various statistical thresholding techniques, such as Bonferroni correction, false discovery rate, concavity of the survival function (Maksymenko et al., 2017), Otsu’s threshold (Chowdhury et al., 2016; Otsu, 1979), and mutual coherence threshold on cortical patches (Krishnaswamy et al., 2017); some of the statistical methods used, are not necessarily designed to be optimal for separating background from signal, and the inverse method itself might have caused the solution to be too smeared for the extent to be meaningful and interpretable. Additionally, some sparse techniques might require a priori knowledge about the solution’s level of sparsity, the number of non-zero elements in the solution vector/matrix, which limits their objectivity. In short, the current MEG source extent estimation is not unique and is largely determined by the researcher’s subjective choice of the thresholding method (or level of sparsity).

In this work, by validating the FAST-IRES on epilepsy patients with MEG recordings, we aim to provide a solution which objectively identifies the location and extent of MEG sources without subjective thresholding. We first evaluate FAST-IRES in computer simulations with realistic head model and spike activity, then validate the capability of FAST-IRES in imaging the epileptogenic tissue from patients’ MEG interictal spikes by comparing the obtained estimates to seizure onset zone (SOZ) determined from invasive iEEG recordings and/or surgical resection volume (whenever available).

## Materials and Methods

### FAST-IRES algorithm

The “fast spatiotemporal iteratively reweighted edge sparsity minimization (FAST-IRES)” algorithm (Sohrabpour et al., 2020) employed in this work is designed to image the location and extent of spatial-temporal neural sources from non-invasive electromagnetic recordings. It first delineates the time-basis function (TBF) of the underlying activity using blind source separation techniques, and then solves for the cortical current density, with an objective function to minimize both the source sparsity and source edge sparsity, while fitting the scalp electromagnetic measurements.

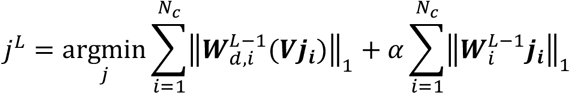

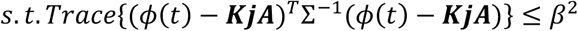

Where Φ(*t*) is the scalp magnetic field measurement (or scalp potential), *K* is the lead-field matrix, *j* is the unknown current density of the brain regions, *A* is the time course activation matrix obtained using blind source separation methods, *β*^2^ is the noise power, Σ is the covariance matrix of the noise to be determined from the baseline activity. 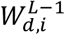 and 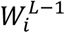 are the weights pertaining to each *j*_*i*_. *V* is the discrete gradient operator, *α* is the hyper-parameter balancing between the two terms of the objective function, and *L* is counting the iteration steps. The algorithm basically, imposes underlying sources to be focally extended; that is, estimates are not to be too focal and not overly diffused, either. The two terms in the objective function, i.e., the source sparsity term and the edge sparsity term, basically ensure that sources retain a balance between becoming too sparse or too dispersed as would be preferred by either term in the objective function. These terms together with the data-fitting constraint, force focally extended solutions and uses a data-driven iterative scheme to penalize locations with smaller amplitudes to quickly converge to an extended source, objectively, without the need for subjective thresholding.

This optimization problem could be solved efficiently and optimally using basic convex optimization tools, as mentioned in the supplementary material in (Sohrabpour et al., 2020). The MATLAB implementation, together with example data are available at https://github.com/bfinl/FAST-IRES.

### Simulation protocol

The default anatomy FSAvg (Fischl et al., 1999) in Brainstorm (Tadel et al., 2011) was used as the head model. The segmented cortex was down-sampled to 30,004 vertices (high-resolution cortex) and 15,002 vertices (low-resolution cortex). A one-layer boundary element method (BEM) model containing the brain was built to calculate the lead-field matrix using OPENMEEG toolbox. The Elekta MEG system with 102 magnetometer and 204 gradiometer sensors was used as the channel configuration. Note that the high-resolution cortex was used for forward modeling and the low-resolution cortex was used for the inverse, to avoid the influence of “inverse crime”. The “inverse crime” happens when the exact model parameters are employed in the forward and inverse modeling in an inverse problem, which might lead to over-optimistic expectations and results (Chávez et al., 2013; Kaipio and Somersalo, 2007). For instance, in source imaging, if the exact same grid and source locations are utilized for the forward and inverse lead-field matrix, the most obvious form of inverse crime has happened!

The procedure for generating simulated MEG signals is depicted in Figure 1A. First, 500 cortical seed locations with a distance (to the closest MEG sensor) less than 60 mm, were randomly selected on the cortex (see supplemental notes for details). Then extended sources were generated using a region-growing approach (Chowdhury et al., 2015) centered at these seed locations, with their radius sampled from a uniform distribution within the 10 mm to 60 mm interval. The source orientation was set to be normal to the cortex. The source activity representing an artificial spike was multiplied by the lead-field matrix to generate the forward solution at sensor space. Lastly, Gaussian white noise of SNR = 5, 10, 20 dB (Liu et al., 2002; Lin et al., 2006; Mattout et al., 2006; Mäkelä et al., 2018; Pantazis and Adler, 2020; Sohrabpour et al., 2020) was added to the sensor space to simulate the noise-contaminated MEG signals.

**Figure 1.**
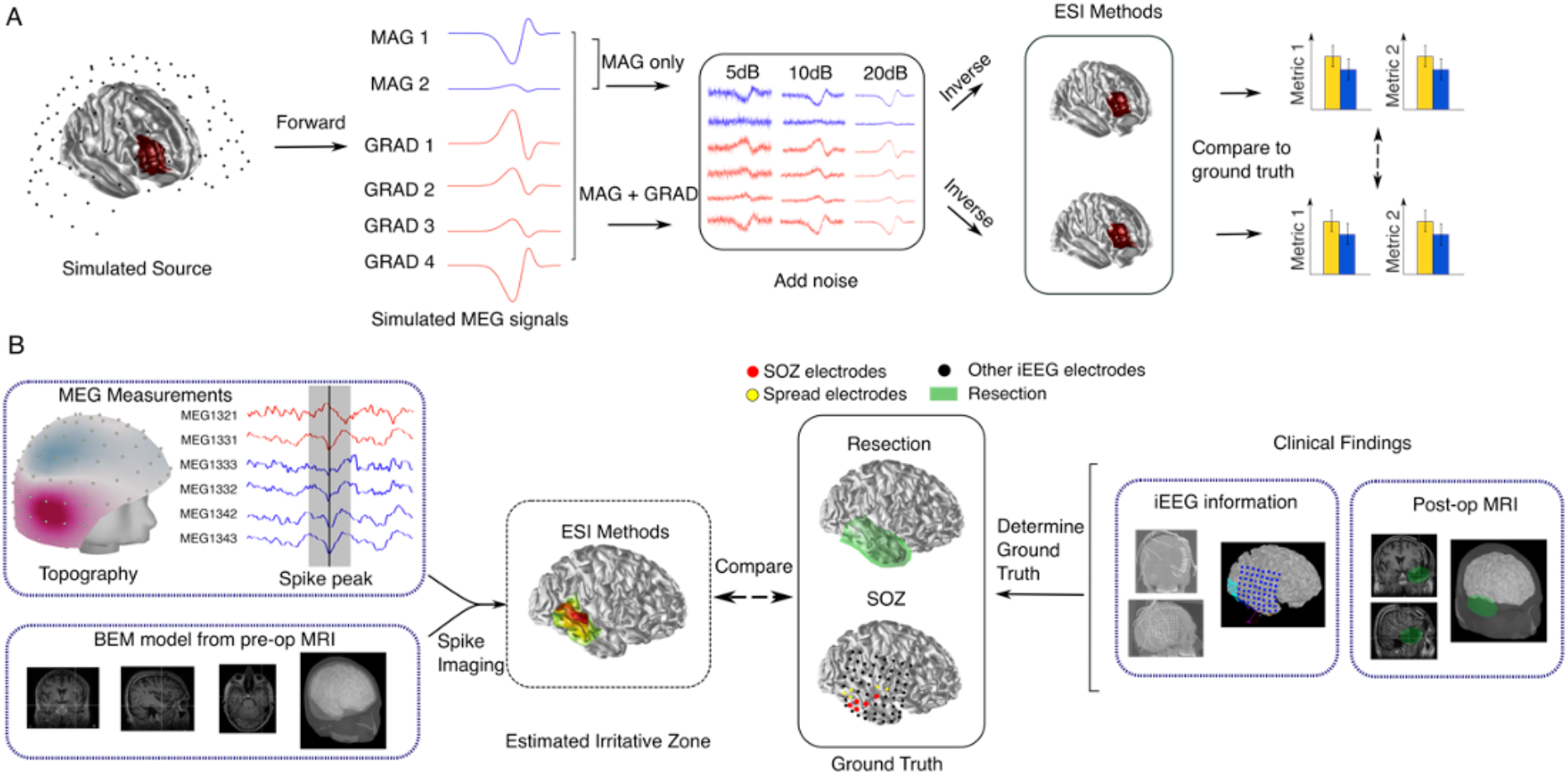
Study design. (A) Simulation protocol. Simulated MEG signals were obtained after choosing the location and extent of the sources. Then Gaussian white noise was added to the clean signal. Different ESI methods (FAST-IRES and LCMV) were performed on the MEG signal to obtain ESI solutions. Finally, performances were compared. (B) Patient analysis protocol. Subject-specific head models were obtained using Brainstorm, and ESI methods were performed to get the solution. This solution was then compared with ground truth, SOZ and resection, which was obtained from clinical iEEG recording reports and post-op MRI.

FAST-IRES performance was compared with LCMV beamformer (Van Veen et al., 1997) as a benchmark ESI algorithms, using the following four performance metrics:

a. Localization error, defined as the distance between the center of the simulated source and the center of mass of the estimated solution.

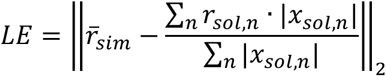 Where 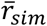 is the center of the simulated source patch, *r*_*sol,n*_ is the n^th^ location at estimated source, and *r*_*sol,n*_ is the ESI solution at n^th^ location.
b. Spatial dispersion, defined as weighted summation of distance between source location and true center location (Hauk et al., 2011; Molins et al., 2008).

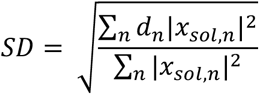 Where *d*_*n*_ is the distance between true center and n^th^ location at estimated source, and *r*_*sol,n*_ is the ESI solution at n^th^ location.
c. Geometric mean of precision and recall. Precision is defined as the ratio of overlapped area between true (simulated) source area and estimated source area, and estimated source area. Recall is defined as the ratio between overlapped are and true (simulated) source area.
d. Pearson’s correlation coefficient between the estimated extent and the simulated (real) extent of simulated sources.

LCMV solutions were then thresholded to give an estimate of the source extent. The threshold was set to be 80% of the neural activity index (NAI) maximum, based on prior studies (Sabeti et al., 2016) and empirical testing (see the supplementary materials).

The ratio between minimum and maximum singular value of the lead-field matrix at each source location was defined as follows(Ahlfors et al., 2010):

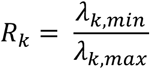

Where *λ*_*k,min*_ and *λ*_*k,max*_ are the minimum and maximum singular value for a *N*_*channel*_ by 3 lead-field matrix at each source location. This ratio is a measure of relative sensitivity to the ‘suppressed’ source orientation.

### Patient analysis protocol

The patients included in this study underwent MEG and intracranial EEG monitoring at the University of Pittsburgh Medical Center (UPMC) as a clinical routine. The data analysis study was approved by and performed in accordance with the regulations of the Institutional Review Board (IRB) of Carnegie Mellon University.

A total of 8 focal drug-resistant epilepsy (DRE) patients were included in this study. The patients were selected based on the following criteria: 1) Patients who underwent pre-surgical evaluation, including MEG recordings and high-resolution MRI; 2) patients who underwent iEEG monitoring and the identified SOZ electrodes were located on the cortical regions (rather than subcortical locations such as amygdala or hippocampus); 3) patients with a clear pattern of interictal spikes; 4) patients who underwent resective surgery and had a postoperative follow-up of at least 12 months. The surgical outcomes were defined based on Engel Surgical Outcome Scale (Engel Jr, 1993), by clinicians. Within the 8 patients, 6 patients were rated as Engel I (free of disabling seizures), and 2 were rated as Engel II (rare disabling seizures).

Each patient underwent a 60-minute 306 channel recording using the Elekta MEG system (Elekta Neuromag, Helsinki, Finland) with 102 magnetometers and 204 planar gradiometers; a 10-minute run was used in this study. The recorded MEG was band-pass filtered between 1-50 Hz with stopband attenuation of 60 dB. For each patient, an individual one-layer boundary element method (BEM) model was constructed from the pre-operational MRI and the sensors were co-registered using anatomical landmarks. Specifically, all surfaces and sensors were first converted from the Elekta/Neuromag coordinate into the CTF coordinate system when imported into Brainstorm software. Then they were co-registered with landmarks such as nasion, left pre-auricular point (LPA), and right pre-auricular point (RPA); digitized head points were also used to refine the registration. The lead-field matrix was then calculated using the OpenMEEG software in MATLAB. The aforementioned preprocessing steps were all conducted with Brainstorm software (Tadel et al., 2011).

For each patient, the MEG waveform was visually inspected to identify the interictal spikes. Two researchers must agree on the identified spikes, for an identified event to be listed as interictal spike. For LCMV, the spikes were averaged before feeding into the algorithm to increase SNR; For FAST-IRES, the blind source separation was performed on the concatenated spikes as part of the preprocessing steps embedded within this algorithm. After that, the algorithm runs the estimated time-basis function (TBF) on averaged spikes after ICA noise removal. For validation, seizure onset electrodes identified from iEEG were extracted and marked by experienced epileptologists in all the patients studied. In 3 patients whose post-operational MRIs were obtained 3-4 months after the resection, the resected volume was extracted and co-registered to the pre-operational MRI and used for comparison.

Similar metrics as in simulation were used in the patient analysis. For patients with SOZ, localization error was defined as the minimum distance between the maximum location of FAST-IRES/LCMV solution, to the closest SOZ electrode. Spatial dispersion was defined as the weighted sum of the minimum distance between each point in the solution to SOZ. For patients with post-op MRI, the localization error was defined as the minimum distance between the maximum location of FAST-IRES/LCMV, to the boundary of resected volume. Note that for the LE calculation, we used the source location with maximum activation rather than center of mass, because for LCMV solutions that are widespread, the center of mass could be deviated from the bulk of activity, making the results not interpretable (see supplementary notes).

Two-sided Wilcoxon rank-sum test was used for statistical comparison between results groups. All the statistical tests were done in MATLAB (version R2019b).

## Results

### FAST-IRES simulation results with MAG group only and MAG+GRAD groups

As shown in Figure 2, under various SNRs (5, 10, 20 dB), FAST-IRES gave robust estimates of the source location and extent. The average LE were 9.93, 9.21 and 8.88 mm, average SD were 4.07, 3.36 and 3.14 mm, and the average geometric mean (of precision and recall - geomean) were 0.63, 0.63 and 0.63 for 5, 10, and 20 dB SNRs, respectively. Statistical difference was found between the LE in the 5 dB 20 dB conditions, the SD of 5 dB and 10 dB, and 5 dB and 20 dB SD (p < 0.05, p < 0.001, p < 0.001, rank-sum test), even though the absolute difference was small. This indicates that in the simulation setting, SNR could influence the FAST-IRES’s performance, but the effect is small. Additionally, FAST-IRES was able to estimate the extent with good accuracy, as reflected by the geometric mean (of precision and recall) values shown in Figure 2A, and the estimated extents versus true extents as plotted in Figure 2C. Figure 2B shows one example of FAST-IRES solution, where the source is at the temporal lobe with different extents, and it could be seen that the estimated extent grows larger as the simulated source’s extent grows larger.

**Figure 2.**
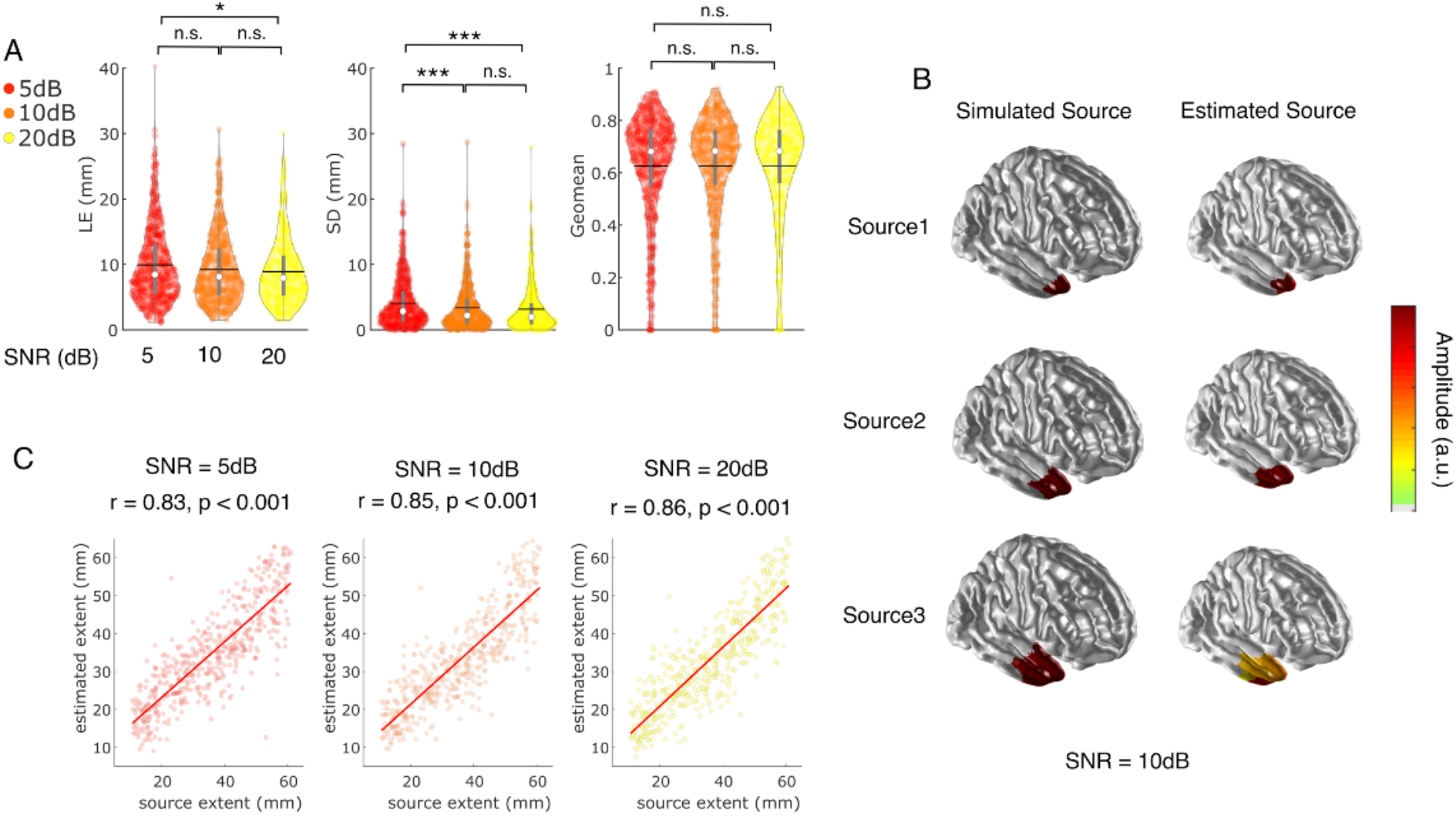
FAST-IRES results under different SNRs. Results are obtained with the MAG sensors only. (A) Localization error (LE), spatial dispersion (SD), and geometric mean of precision and recall (geomean) under different SNR scenarios. Statistical comparison results are obtained using Wilcoxon rank sum test, *: p < 0.05, **: p < 0.01, ***: p < 0.001). In each violin plot, the black horizontal bar indicates the mean, the white dot indicates the median, the grey vertical bar indicates the interquartile range (the difference between 75th and 25th percentiles), and the envelope indicates the distribution of the data. (B) FAST-IRES example estimation solutions. Source 1 to 3 are all at the same location but with different extents. The color bar indicates the amplitude of the solution. (C) Correlation between the simulated extent and source extent, the red line indicates the least square regression line.

Figure 3 shows the results using the MAG+GRAD configuration. The average LE for 5, 10, and 20 dB SNRs are, respectively, 8.58, 7.68 and 7.65 mm, with the average SD being 3.02, 2.93 and 2.93 mm, and the average geomean is 0.63, 0.64 and 0.64. Significant difference was found between 5dB and 10dB, s5dB and 20dB for LE (p < 0.001, rank-sum test). Additionally, Figure 3D shows the comparison between results with MAG and MAG+GRAD, and a statistically significant difference could be found between these metrics in certain SNRs (also see Supp Fig1). This particular result is not surprising as adding the gradiometer could potentially offer more information regarding the source of activity. However, our results suggest that such information might not necessarily lead to better extent estimation of underlying sources, as reflected by the insignificant difference between the geomean of sources under these different configurations.

**Figure 3.**
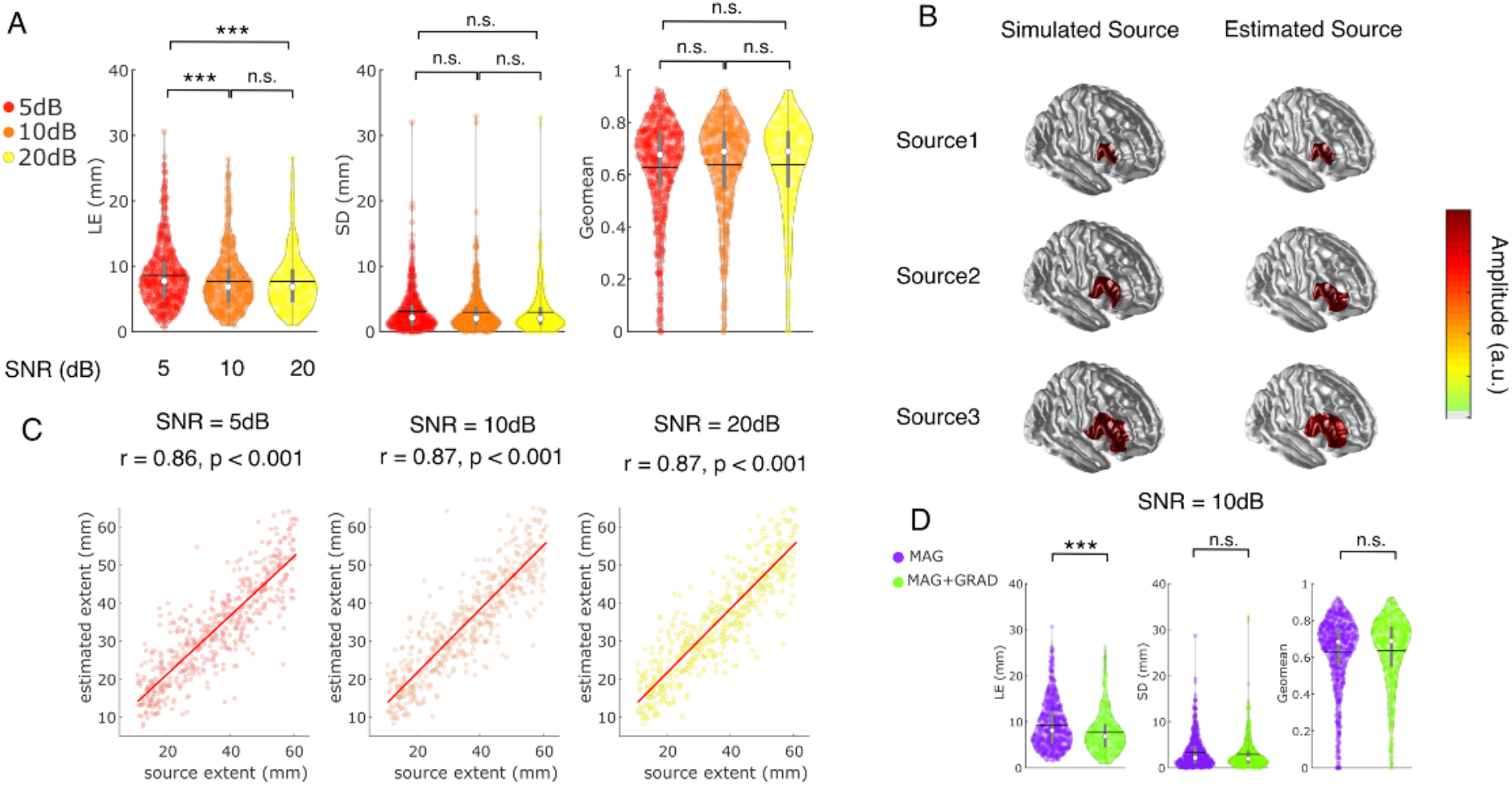
FAST-IRES results under different SNRs. Results are obtained with MAG+GRAD sensors. Panel A-C are similar to Figure 2 A-C. In each violin plot, the black horizontal bar indicates the mean, the white dot indicates the median, the grey vertical bar indicates the interquartile range (the difference between 75th and 25th percentiles), the envelope indicates the distribution of the data, the red line indicates the least square regression line, and the color bar indicates the amplitude of the solution. (D) Comparison between MAG and MAG+GRAD in 10 dB scenario.

### Influence of source depth and location on the performance of FAST-IRES

Figure 4 shows how FAST-IRES performance varies across different locations and depths, on the cortex.

**Figure 4.**
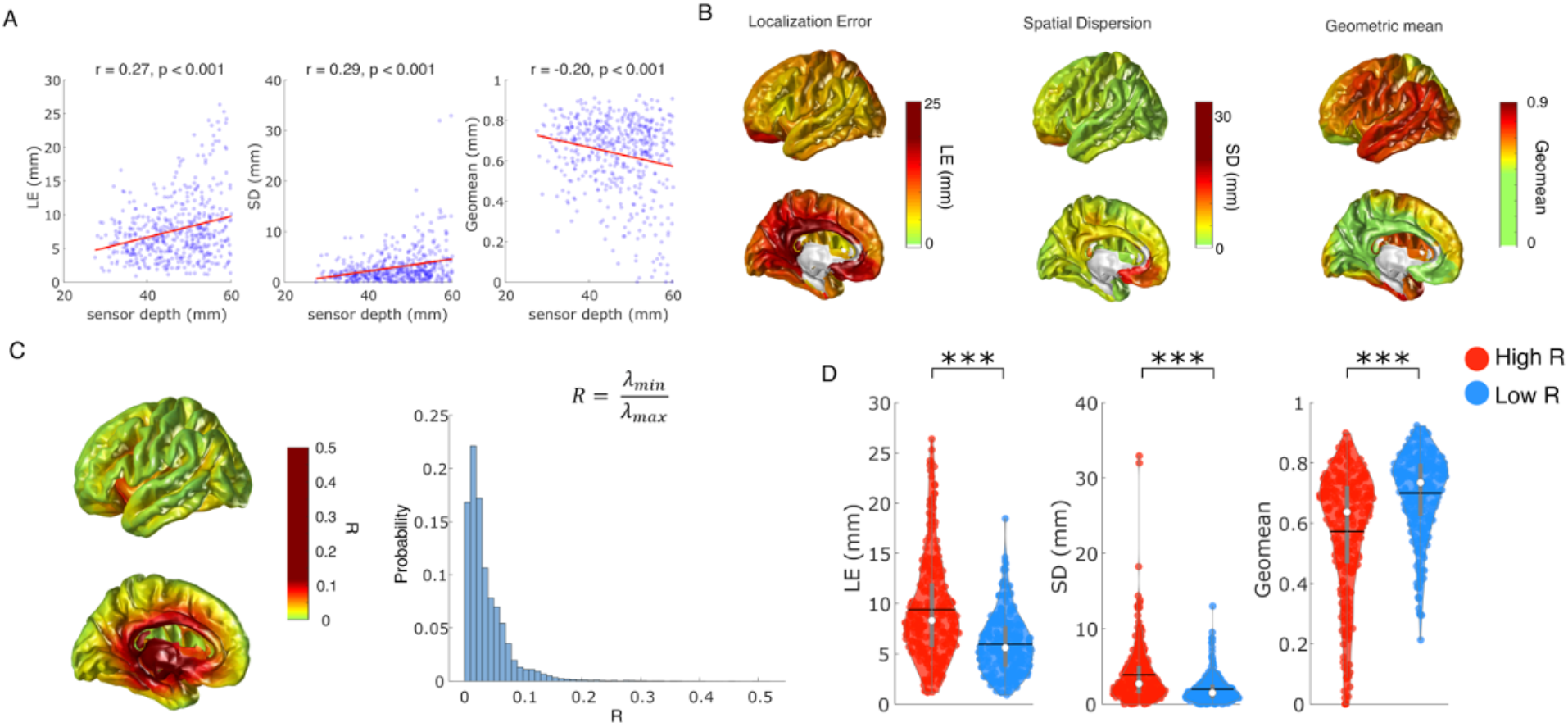
FAST-IRES performance at different locations and depth. (A) Correlation between the source depth (defined with respect to distance from closest MEG sensor), and localization error (LE), spatial dispersion (SD), and the geometric mean of precision and recall (geomean). A significant correlation could be found. (B) distribution of these performance metrics over the cortex. Notice that the color bar scheme is different among these performance metrics, to illustrate the best contrast within the plot. (C) Ratio (*R*) of between minimum and maximum singular value (*λ*_*min*_ and *λ*_*max*_) of the lead-field matrix at each source location, data is shown as colormap on the cortex and histogram. (D) Performance comparison between high R and low R sources.

Figure 4 A demonstrates the correlation between source depth and LE, SD and geomean, in the MAG+GRAD, 10 dB noise condition. The correlations are 0.27, 0.29, and −0.20, respectively, with p < 0.001. As sources go deeper, the LE and SD increases while geomean decreases, indicates that deeper sources are more difficult to retrieve than superficial sources. Figure 4 B shows the distribution of these performance metrics over the cortex. For each extended source, all cortical points belonging to this source region is given a value equal to the performance; for example, if the localization error for this source is 10mm, then all cortical points in this extended source is given a value of 10 mm. If one cortical point belongs to multiple sources, then the value at this point is the averaged over these sources. The observation is consistent with the numerical evaluation that superficial sources typically have better performance.

Figure 4 C describes the distribution of ratio (R) between minimum and maximum singular value of the lead-field matrix at each source location, used previously (Ahlfors et al., 2010) to indicate the sensitivity to the suppressed source orientation. A low value or R indicates that there is more difference between the directions corresponding to max and min lead-field gain. The appearance and distribution of R over the cortex is consistent with Ahlfors and colleagues’ work. A certain trend could be observed that high R typically occurs in the sulci patch and low R typically occurs in the gyral patch. Next, we classified sources into high R and low R, in Figure 4 D it is found that high R sources have larger LE, SD and lower geomean, indicating a worse performance. This result indicates that in general low R sources will have better performance.

### FAST-IRES simulation results compared with other ESI methods

Supp Fig 2C, D shows the results of FAST-IRES compared with LCMV methods. A statistically significant difference was found between FAST-IRES and LCMV (p < 0.001, rank-sum test). Supp Fig 2B shows two examples of the solution, which demonstrates that FAST-IRES not only accurately estimate the location, but also the extent of the simulated sources. The results also indicate that with the varying source extent model, LCMV with an arbitrary threshold failed to capture the extent of the simulated source well, even though we carefully chose the threshold value.

### Evaluation of FAST-IRES in drug-resistant epilepsy patients

Figure 5 shows the evaluation results in a group of drug-resistant epilepsy patients. From the examples in Figure 5A-C, it could be observed that the FAST-IRES solution is in general pointing to the ground truth. On the other hand, LCMV solutions are typically widespread (see Supp Fig 3). To illustrate the overlap between SOZ and IZ electrodes. On average 62% of the SOZ electrodes also belong to IZ electrodes, indicating a certain amount of overlap between the SOZ and IZ. For the 6 patients that are seizure-free (Engel I), the averaged LE compared to SOZ for FAST-IRES and LCMV are, respectively, 15.5, 15.7 mm, the average SDs are 20.8, 35.9 mm. For all the 8 patients, the averaged LEs are 17.0, 19.3 mm, the averaged SDs are 21.3, 38.9 mm, the difference was significant between FAST-IRES SD and LCMV SD (p < 0.05, rank-sum test). For the results (when using resection as ground truth), the averaged LEs for FAST-IRES and LCMV are, respectively, 12.4, 23.4 mm, the averaged SDs are 20.6, 29.1 mm. We also calculated these performance metrics with respect to the IZ electrodes. The IZ is defined as the electrode locations where spikes were recorded in the clinical report, these electrodes could be further classified as primary IZ with most frequent spikes, and other IZ with occasional spikes (Figure 5). The average LE and SD are 16.1 mm and 19.0 mm. One potential explanation for the performance being better with respect to IZ is that IZ is typically larger than SOZ, which LE and SD are in favor of. Based on the above results, a similar observation could be made that FAST-IRES is performing reasonably well, able to estimate both location and extent of the inter-ictal sources.

**Figure 5.**
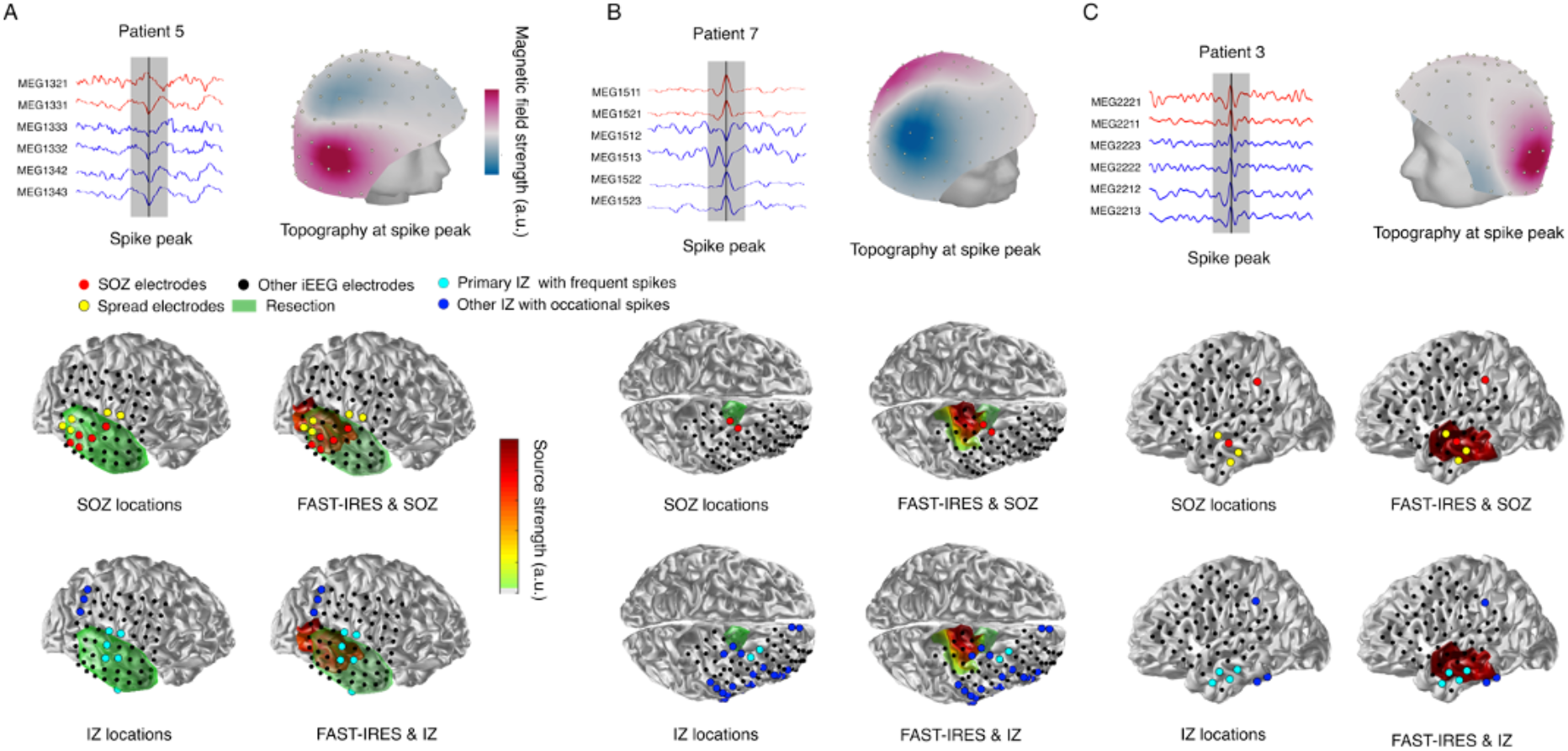
Patient analysis results. (A, B) ESI results in example patients, compared with SOZ (red and yellow) and resection (green), IZ electrodes are also shown as reference (based on the number of occurrences of spikes, electrodes are further classified into major IZ (cyan) and other IZ (blue)). (C) Example patient compared with SOZ only (since this patients’ post-operative MRI is not available), IZ electrodes are also shown.

## Discussion

Imaging spatiotemporally distributed epileptiform sources from MEG measurements has been an important aspect in pre-surgical evaluation. In this work, through rigorous simulations and validation in 8 focal epilepsy patients, we demonstrated the capability of the FAST-IRES algorithm to estimate the location and extent of the underlying epilepsy sources from MEG measurements, objectively and noninvasively.

The source extent is of significance as the ability to distinguish between the relevant epileptiform activity and background activity is a necessary requirement for determining surgical margin, which offers valuable information for pre-surgery planning. Previous work that estimated the extent from MEG measurements implemented various thresholding techniques on the solution (Bouet et al., 2012; Attal and Schwartz, 2013; Krishnaswamy et al., 2017; Pellegrino et al., 2018), but a major shortcoming is that the extent estimation by means of applying a threshold to the solution, is subjected to the researcher’s choice. In our study, the choice of LCMV threshold was based on both prior literature and empirical testing, aiming to give the optimal geometric mean for comparison. Although other more advanced statistical thresholding methods exist, such as Otsu’s threshold (Chowdhury et al., 2016), Bonferroni correction, false discovery rate and concavity of the survival function (Maksymenko et al., 2017), to the best of our knowledge the empirical thresholding is still the most common approach for extent estimation in these algorithms (Palmero-Soler et al., 2007; de Gooijer-van de Groep et al., 2012; Attal and Schwartz, 2013; Sun and Kobayashi, 2017). Nevertheless, FAST-IRES still outperformed the benchmark method with a lower spatial dispersion and a higher geometric mean of precision and recall. Our results indicate that FAST-IRES is able to capture the correct extent of the underlying sources, while LCMV were not able to provide accurate extent estimates, in spite of careful thresholding.

In the simulation, for all the employed performance metrics, namely LE, SD, geometric mean and correlation between simulation extent and estimated extent, FAST-IRES results remain largely consistent between different SNR scenarios. This observation is in agreement with our prior work on EEG (Sohrabpour et al., 2020) and demonstrates the robustness of our approach with respect to noise level. One reason for this observation, is that FAST-IRES incorporates an adaptive noise estimation process, which controls the size of hyper-ellipsoid constraining the output power (Sohrabpour et al., 2016a). This property enables the adaptability of FAST-IRES in real MEG analysis, as varying levels of noise is typically encountered in practice due to different equipment and recording environment. On the other hand, there is prior work comparing the performance of EEG and MEG, showing that MEG source estimates are more superficial than EEG (Leijten et al., 2003), making them slightly disadvantageous than EEG in imaging deeper tissue, which is consistent with the fact that MEG is more sensitive to cortical sources (Baillet, 2017), MEG source imaging was also shown to yield better concordance and distance from the clinical SOZ than EEG source imaging (Pellegrino et al., 2018). Nevertheless, it is also notable that in majority of these research, MEG usually have more sensors than EEG electrodes, which may contribute to the performance as well.

One unique aspect of MEG modality compared to EEG is that MEG has different types of sensors, MAG and GRAD, and various channel configurations have been used in prior works to perform source imaging (Henson et al., 2009; Wens et al., 2014; García-Pacios et al., 2015; Hillebrand et al., 2016; Garcés et al., 2017). In terms of FAST-IRES, in general we found that MAG+GRAD and MAG group have comparable results, both in simulation and in patient analysis. This observation indicates that MAG group has already captured most of the information regarding the source location and extent, and could serve as a reference when one is considering sacrificing computation time for better source imaging accuracy using more MEG channels, in FAST-IRES. Nevertheless, it should be noted that such a conclusion is drawn under the specific MEG system (Elekta) we used. Other different MEG systems, such as CTF (Fife et al., 1999), have different sensor configurations, and it remains to be investigated if such a conclusion is still applicable in other configurations.

One limitation of our study is that the experimental design and conclusions were based on analyzing cortical sources. This is largely due to the fact that MEG is less sensitive to deep cortical sources and subcortical sources (Agirre-Arrizubieta et al., 2009; Attal et al., 2009; Ahlfors et al., 2010; Wennberg et al., 2011; Attal and Schwartz, 2013). Deep source localization has long been considered a difficult problem (Hillebrand and Barnes, 2002; Attal et al., 2009), especially given the need for the development of radically new algorithms capable of achieving such a goal (Krishnaswamy et al., 2017; Bénar et al., 2021). Recent attempts have been made to address the issue, but such studies usually require the aid of simultaneous intracranial recordings (Juárez-Martinez et al., 2018; Pizzo et al., 2019; Pellegrino et al., 2020) or prior knowledge of the activation patterns (Krishnaswamy et al., 2017) while large-scale validation is still not available. On the other hand, FAST-IRES is currently implemented as a cortex-based algorithm, but it can, in principle, be expanded to volume-based sources to better estimate subcortical regions. That said, further exploration must be done in the future to adapt our proposed source imaging algorithm for subcortical region and deep structure localization.

In conclusion, we showed that FAST-IRES can be generalized to MEG source imaging by demonstrating its ability to image the location and extent of underlying epilepsy sources from MEG measurements, both in simulations and in a clinical study of 8 drug-resistant epilepsy patients. Our results indicate FAST-IRES may aid the pre-surgical planning in drug-resistant epilepsy patients by providing MEG source imaging solutions with a more objective and robust estimation of both the location and extent of epileptogenic tissues.

## Acknowledgement

This work was supported in part by National Institutes of Health grants NS096761, EB021027, MH114233, EB029354, and AT009263. We are grateful to Drs. Alexandra Urban, Mark Richardson and Vasileios Kokkinos for their respective clinical roles in evaluating and treating surgically included patients, and Zhengxiang Cai for helpful discussions on data analysis. We thank participating patients and their families whose involvement and sacrifice made this work possible. We also acknowledge selfless dedication and invaluable efforts of the University of Pittsburgh Comprehensive Epilepsy Center (UPCEC) team and particularly the staff of the UPMC Presbyterian University Hospital (PUH) Epilepsy Monitoring Unit (EMU) led by Cheryl Plummer, BS, R. EEG T., NA-CLTM, FASET.

## Notes

### Competing Interest Statement

The authors have declared no competing interest.

